# High genomic plasticity and unique features of *Xanthomonas translucens* pv. *graminis* revealed through comparative analysis of complete genome sequences

**DOI:** 10.1101/2023.06.29.547023

**Authors:** Florian Goettelmann, Ralf Koebnik, Veronica Roman-Reyna, Bruno Studer, Roland Kölliker

**Author notes:** Correspondence: Roland Kölliker.

## Abstract

**Background:** *Xanthomonas translucens* pv. *graminis* (*Xtg*) is a major bacterial pathogen of economically important forage grasses, causing severe yield losses. So far, genomic resources for this pathovar consisted mostly of draft genome sequences, and only one complete genome sequence was available, preventing comprehensive comparative genomic analyses. Such comparative analyses are essential in understanding the mechanisms involved in the virulence of pathogens and to identify virulence factors involved in pathogenicity.

**Results:** In this study, we produced high-quality, complete genome sequences of four strains of *Xtg*, complementing the recently obtained complete genome sequence of the *Xtg* pathotype strain. These genomic resources allowed for a comprehensive comparative analysis, which revealed a high genomic plasticity with many chromosomal rearrangements, although the strains were highly related, with 99.9 to 100% average nucleotide identity. A high number of transposases were exclusively found in *Xtg* and corresponded to 413 to 457 insertion/excision transposable elements per strain. These mobile genetic elements are likely to be involved in the observed genomic plasticity and may play an important role in the adaptation of *Xtg*. The pathovar was found to lack a type IV secretion system, and it possessed the smallest set of type III effectors in the species. However, three XopE and XopX family effectors were found, while in the other pathovars of the species two or less were present. Additional genes that were specific to the pathovar were identified, including a unique set of minor pilins of the type IV pilus, 17 TonB-dependent receptors (TBDRs), and 11 degradative enzymes.

**Conclusion:** These results suggest a high adaptability of *Xtg*, conferred by the abundance of mobile genetic elements, which may have led to the loss of many features. Conserved features that were specific to *Xtg* were identified, and further investigation will help to determine genes that are essential to pathogenicity and host adaptation of *Xtg*.

## 1 Introduction

Bacterial wilt of forage grasses, caused by *Xanthomonas translucens* pv. *graminis* (*Xtg*), is one of the main diseases of forage grasses in temperate grasslands (1). *Xtg* is a member of the recently-defined clade Xt-III of *X. translucens*, which also contains four other forage grass infecting pathovars (pv. *arrhenatheri*, *phlei*, *phleipratensis* and *poae*) and is distinguished from clades Xt-I and Xt-II, which contain mainly cereal-infecting pathovars such as pv. *translucens* and *cerealis*, respectively (2,3). While most *X. translucens* pathovars are generally restricted to a small host range, *Xtg* is able to infect many grass genera, including *Agrostis*, *Alopecurus*, *Dactylis*, *Deschampsia*, *Festuca*, *Lolium*, *Phalaris*, *Phleum*, *Poa*, and *Trisetum* (1,4). It is also the most widespread, being found in most of Europe, America, and New Zealand (4–6).

The bacteria are mainly spread by contaminated mowing tools and penetrate via plant wounds. They then spread in the xylem, resulting in a disruption of the flow of water and nutrients (7). The disease affects mainly Italian ryegrass (*Lolium multiflorum* Lam.), which is cut several times during the growing season for harvesting. Infection by the pathogen can lead to serious yield losses, estimated to reach up to 40% under unfavourable conditions (8). To control the disease in an efficient manner, it is essential to understand the mechanisms involved in the pathogenicity of *Xtg*.

In plant pathogenic bacteria, successful infection of the plant host generally involves virulence factors that allow the bacteria to adhere to the plant surface, invade the host tissue, acquire nutrients, and suppress plant defence mechanisms. In Xanthomonads, some of the most important virulence factors are effectors, in particular type III effectors (T3E), that are secreted into the plant host cells by the type III secretion system (T3SS). These T3E play a major role in pathogenicity by specifically targeting different pathways in the host cell. A total of 53 classes of T3E have been defined, generally referred to as “*Xanthomonas* outer proteins” (Xop) (9). Other T3E can cause a hypersensitive response in their plant host and are referred to by this characteristic, like AvrBs1 and AvrBs2. Additionally, transcription activator-like effectors (TALE) are specific T3E that consist of repetitive sequences that allow them to bind to specific nucleotide sequences in the host genome and manipulate the expression of specific genes (10,11). However, while TALE are characteristic to the *Xanthomonas* genus, no TALE have ever been identified in *Xtg* (3,12).

Other secretion systems also play an important role in the infection of *Xanthomonas* spp., including the type II, IV and VI secretion systems (T2SS, T4SS and T6SS, respectively). The T2SS is mostly involved in the secretion of plant cell wall degradative enzymes in the apoplast, allowing the bacteria to invade the plant host, as well as acquire the resulting degradation products as a source of nutrients (13). Two types of T2SS can be found in *Xanthomonas* spp., the *xps* T2SS, which is conserved across the genus, and the *xcs* T2SS, which is only found in some species (14). To date, no *xcs* T2SS was found in *X. translucens* (3).

The T4SS and T6SS have a similar function and are involved in defence against microbial predators and competition with other microorganisms (15,16). Three subtypes of the T6SS have been found in *Xanthomonas* spp., subtypes 1, 3 and 4, and subtype 3 is further divided into subgroups 3*, 3** and 3***. Previously, no T4SS has been identified in *Xtg*, and T6SS genes were identified only in strains Xtg2, Xtg9, Xtg10 and NCPPB 3709 (3,17).

To reliably identify virulence factors in plant pathogenic bacteria, complete and high-quality genome sequences are essential, as they provide an accurate representation of the genes present in each strain. Such genome sequences enable comparative genomics, which can reveal conserved or divergent genes of a pathovar and help define genes that are crucial to pathogenicity. Previously, draft genome sequences of *Xtg* strains CFBP 2053, ICMP 6431, NCPPB 3709, Xtg2, Xtg9 and Xtg29 allowed a first comparative analysis, which revealed specific features of the pathovar, such as the lack of a flagellum and a distinct type IV pilus (17). Hypothetical proteins specific to *Xtg* were also identified, including genes with predicted functions in nutrient acquisition, regulatory mechanisms, virulence, adhesion and motility. Effector proteins that were specific to *Xtg* included a YopT-like cysteine protease and a XopJ class effector protein, which were tested for their role in virulence. However, single and double knock-out mutants for these genes showed no significant difference in virulence with the wild type (17).

At that time, genomic resources for *X. translucens* pathovars consisted of highly fragmented and incomplete draft genome sequences based on short read sequencing, which might have prevented the identification of important genes. Furthermore, these genome assemblies fail to assemble complex repetitive sequences such as TALEs, which may have hindered their identification. Long read sequencing technologies now allow to assemble complete bacterial genome sequences, allowing for a comprehensive overview of the gene content within these genomes, including TALEs (18). Recently, high-quality, complete genome sequences were produced for all pathotype strains of *X. translucens*, which allowed for a more comprehensive comparative analysis (3). This revealed that the *Xtg* pathotype strain LMG 726 lacked many virulence features, such as a T4SS and T6SS, and had the smallest set of T3E. However, the inclusion of only one strain of *Xtg* did not allow for an in-depth comparative analysis. In this study, we aimed at producing high-quality complete genome sequences for four additional strains of *Xtg,* perform a comprehensive comparative genomic analysis of the pathovar, and identify unique features that could play a role in its virulence and host specificity.

## 2 Materials and methods

### 2.1 Bacterial strains, growth conditions and DNA extraction

The *Xtg* strains NCPPB 3709, Xtg2, Xtg9 and Xtg29 were selected to be sequenced in this study (Table 1). These strains were grown at 28°C on YDC agar medium (2% dextrose, 1% yeast extract, 2% CaCO3, 1.5% agar) for 48 h. Bacteria were then dissolved in 10 mL washing buffer (50 mM TRIS-HCl pH 8.0, 50 mM EDTA pH 8.0, 150 mM NaCl). Genomic DNA was then extracted with the NucleoSpin® Microbial DNA kit (Macherey Nagel, Duren, Germany), according to the manufacturer’s recommendations.

**Table 1.** *Xanthomonas translucens* strains used for comparative genomic analysis and characteristics of the genome assemblies. The pv. *translucens* and pv. *cerealis* pathotype strains DSM 18974 and CFBP 2541 were included as representatives of clades Xt-I and Xt-II, respectively.

| Pathovar | Strain<sup>1</sup> | Country | Isolated from | Origin<sup>2</sup> | Coverage | Chromosome length (bp) | Plasmid length (bp) | Accession | Reference |
| --- | --- | --- | --- | --- | --- | --- | --- | --- | --- |
| *graminis* | NCPPB 3709 | Norway | *Lolium multiflorum* | NCPPB | 89 | 4,732,427 | 44,971 | CP118161-CP118162 | This study |
|  | Xtg2 | Switzerland | *Lolium multiflorum* | (40) | 78 | 4,722,726 | NA | CP076253 | This study |
|  | Xtg9 | Switzerland | *Lolium multiflorum* | (40) | 84 | 4,734,182 | NA | CP076252 | This study |
|  | Xtg29 | Switzerland | *Lolium multiflorum* | (40) | 64 | 4,673,077 | NA | CP076257 | This study |
|  | LMG 726 | Switzerland | *Lolium multiflorum* | BCCM | 81 | 4,678,781 | NA | CP076254 | (3) |
| *arrhenatheri* | LMG 727 | Switzerland | *Arrhenatherum elatius* | BCCM | 103 | 4,843,101 | NA | CP086333 | (3) |
| *poae* | LMG 728 | Switzerland | *Poa trivialis* | BCCM | 86 | 4,792,655 | NA | CP076250 | (3) |
| *phlei* | LMG 730 | Norway | *Phleum pratense* | BCCM | 87 | 4,569,024 | NA | CP076251 | (3) |
| *phleipratensis* | LMG 843 | USA | *Phleum pratense* | BCCM | 121 | 4,902,099 | NA | CP086332 | (3) |
| *cerealis* | CFBP 2541 | USA | *Bromus inermis* | CFBP | 404 | 4,504,942 | NA | CP074364 | (3) |
| *translucens* | DSM 18974 | USA | *Hordeum vulgare* | DSMZ | 223 | 4,715,357 | NA | LT604072 | (41) |

<sup>1</sup>Pathotype strains are marked in bold.
<sup>2</sup>NCPPB: National Collection of Plant Pathogenic Bacteria (Fera Science Ltd., Sand Hutton, United Kingdom); BCCM: Belgian Coordinated Collections of Microorganisms; CFBP: Collection of Plant Pathogenic Bacteria; DSMZ: German Collection of Microorganisms and Cell Cultures.
<sup>3</sup>TBDRs: TonB-dependent receptors

### 2.2 Sequencing, genome assembly and annotation

Library preparation and DNA sequencing was performed at the Functional Genomics Center Zurich. Libraries were prepared and multiplexed with the PacBio SMRTbell® Express Template Prep Kit 2.0 (PacBio, Menlo Park, CA, USA) according to the published protocol (19). Libraries were then sequenced on the PacBio Sequel II platform.

Genomic sequences were *de novo* assembled with flye 2.7 for the strains Xtg2, Xtg9 and Xtg29 using the “--plasmids --iterations 2” parameters, and canu for NCPPB 3709 using default parameters (20,21). The assembled circular chromosomes were rotated to start with the *dnaA* gene sequence for comparability. Genomes were functionally annotated with Prokka 1.14.6 (22).

### 2.3 Phylogeny and comparative analysis

To determine phylogeny within clade Xt-III, we used all available complete genome sequences of the clade, as well as the genome sequences of pv. *translucens* strain DSM 18974 and pv. *cerealis* strain CFBP 2541 as representatives of clades Xt-I and Xt-II, respectively (Table 1). Average nucleotide identity (ANI) was calculated using pyani 0.2.12 with default parameters (23), and a phylogeny dendrogram was constructed using Ward’s hierarchical clustering method in R version 4.2.0 (24). To identify genomic rearrangements and conserved genomic regions within *Xtg*, the genomic structure of the five *Xtg* genome sequences were compared with Mauve v20150226 (25).

For a comparative analysis of the gene content of *Xtg* compared to other *X. translucens* strains, all currently available complete genome sequences of the *X. translucens* species were retrieved and annotated with Prokka (Table 1, Table S1). The gene content within the *X. translucens* species was compared with Roary 3.13.0 using default parameters, and singletons, genes that were exclusive to *Xtg* strains, were defined (26). The pangenome and the singleton contents of *Xtg* strains were visualized in Venn diagrams with jvenn (27). Clusters of orthologous groups (COG)-based functional characterization of the gene content of *Xtg* was performed using eggNOG-mapper v2 (28,29). As the COG analysis revealed a high number of transposable elements in *Xtg*, these were further characterized using the mobileOG-db tool in the Proksee online software, with default parameters (30,31). Additionally, putative horizontal transfer events were identified using the Alien Hunter tool in Proksee (32). The presence of signal peptides or non-classical secretion were predicted using SignalP 6.0 and SecretomeP 2.0, respectively (33,34).

Minor pilins identified in the singleton analysis were investigated with InterProScan by identifying the characteristic pilin N-terminal transmembrane domain, as well as with the PilFind online tool (35,36). To compare the two filamentous hemagglutinin singleton genes identified with similar previously-identified genes, the nucleotide sequence of these genes in the Xtg29 draft genome sequence in which they were identified was extracted (17). These sequences were then used as query in a BLASTn analysis using the nucleotide sequences of the genes identified in this study or the whole genome sequences of *Xtg* strains as subject. The DNA sequences of these gene clusters were compared with Clinker v0.0.27 (37).

Presence of T2SS, T3SS, T4SS and type III effectors was determined as detailed by Goettelmann et al. (3). Presence of T6SS was determined using SecReT6 3.0 with default settings (38). The presence of TALEs was determined using AnnoTALE 1.5 (39).

## 3 Results

### 3.1 Genome assembly

Each strain was assembled to one circular complete chromosome. The chromosome sequences were of comparable size, ranging from 4,673,077 bp for Xtg29 to 4,734,182 bp for Xtg9. The sequence coverage ranged from 64- to 89-fold for all the sequenced strains (Table 1). Additionally, a circular contig sequence of 44,971 bp was assembled in strain NCPPB 3709. A comparison by BLASTn to the NCBI non-redundant nucleotide sequences database showed homology to plasmid sequences. This contig was thus considered to represent a plasmid of this strain.

### 3.2 Phylogeny and comparative analysis

Phylogeny based on ANI revealed that all strains of *Xtg* were very similar, with 99.9% to 100% ANI (Figure 1). Comparison of the genomic structure within *Xtg* showed 65 to 124 locally collinear blocks (LCB), genomic regions that show no rearrangements between the compared strains (Figure 2). This showed that although *Xtg* strains are very close genetically, a high level of rearrangements can be seen within the pathovar.

**Figure 1.**
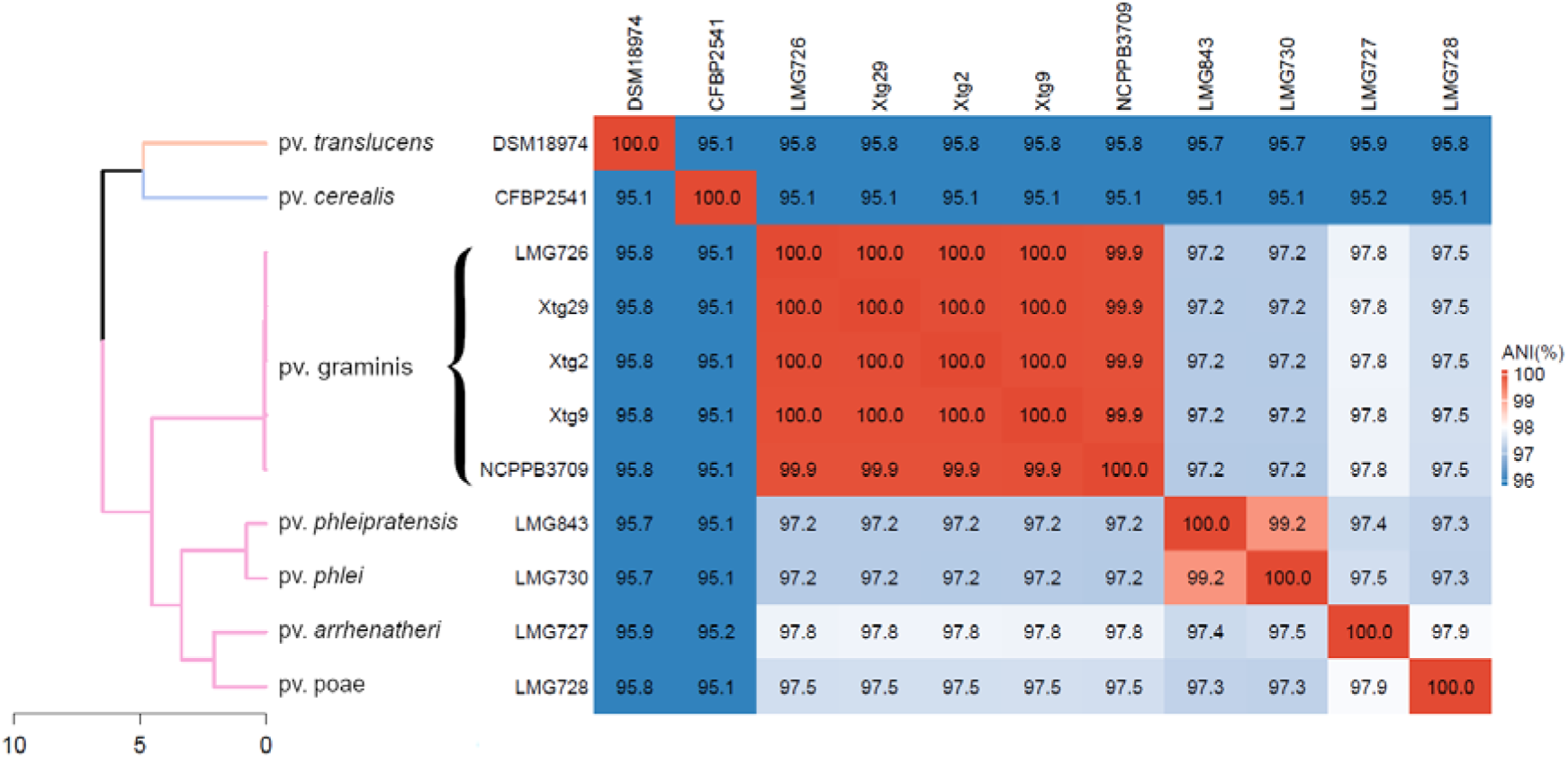
Average nucleotide identity (ANI)-based phylogeny of *Xanthomonas translucens* clade Xt-III, including all complete genome sequences currently available for the clade, as well as DSM 18974 and CFBP 2541 as representatives of clades Xt-I and Xt-II, respectively. Phylogeny was constructed with Ward’s hierarchical clustering method. ANI is depicted as a gradient from blue (< 96%) to white (98%) to red (100%). Distance in the dendrogram represents dissimilarity between nodes. Orange: clade Xt-I, blue: clade Xt-II, pink: clade Xt-III.

**Figure 2.**
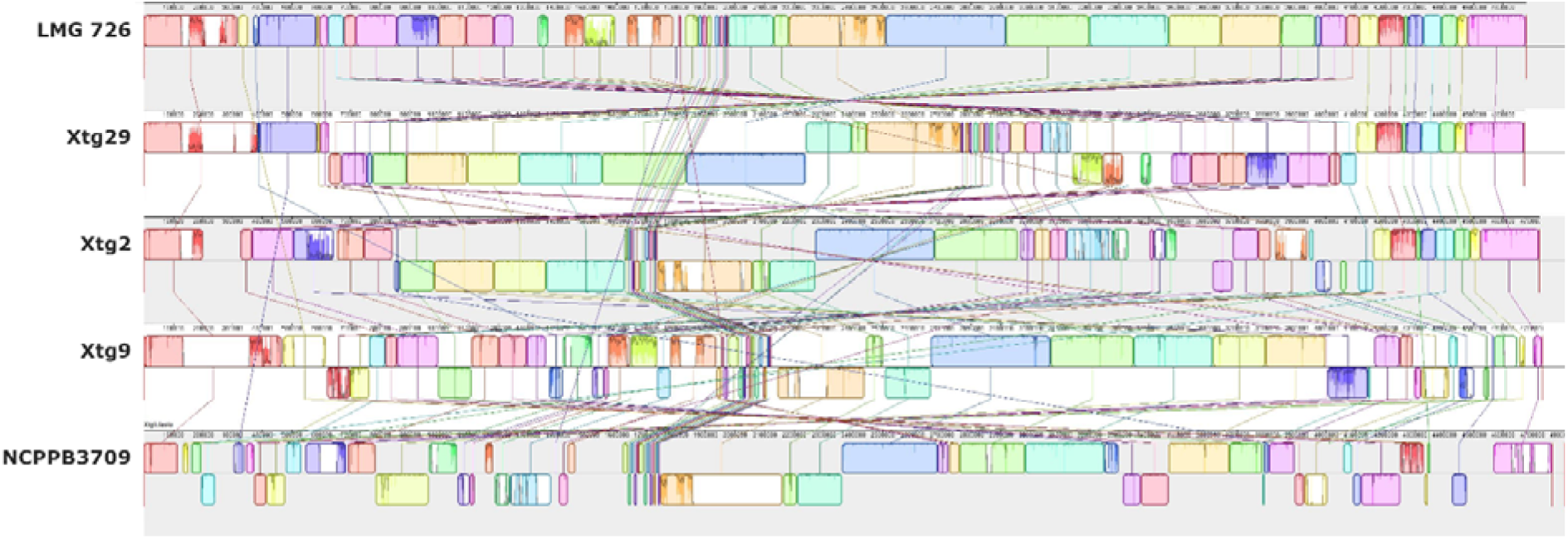
Pairwise comparisons of the chromosome structure within *Xanthomonas translucens* pv. *graminis* using all five currently available complete genome sequences of the pathovar. The LMG 726 genome sequence at the top was used as a reference. Colours represent locally colinear blocks (LCB), regions that show no rearrangement across all the compared genome sequences. Synteny is displayed by a line that links orthologous LCBs. LCBs found on the bottom part represent regions that are found in reverse orientation compared to the reference.

The number of genes found in each strain of *Xtg* was similar, ranging from 4,408 in Xtg9 to 4,445 in NCPPB 3709 (Table 1). The pangenome of the five *Xtg* strains used in this study consisted of 4,931 genes, while the core genome consisted of 3,739 genes (Fig. 3A). Moreover, a total of 1,409 singletons, genes that were not found in any other pathovar of the *X. translucens* species, were identified, with 536 being shared by all *Xtg* strains (Fig. 3B). A total of 433 singletons were found to be strain-specific, while 131 were shared by two strains, 164 were shared by three strains, and 145 were shared by four strains. The total number of singletons per strain ranged from 856 in Xtg9 to 922 in Xtg29 (Table 2).

**Figure 3.**
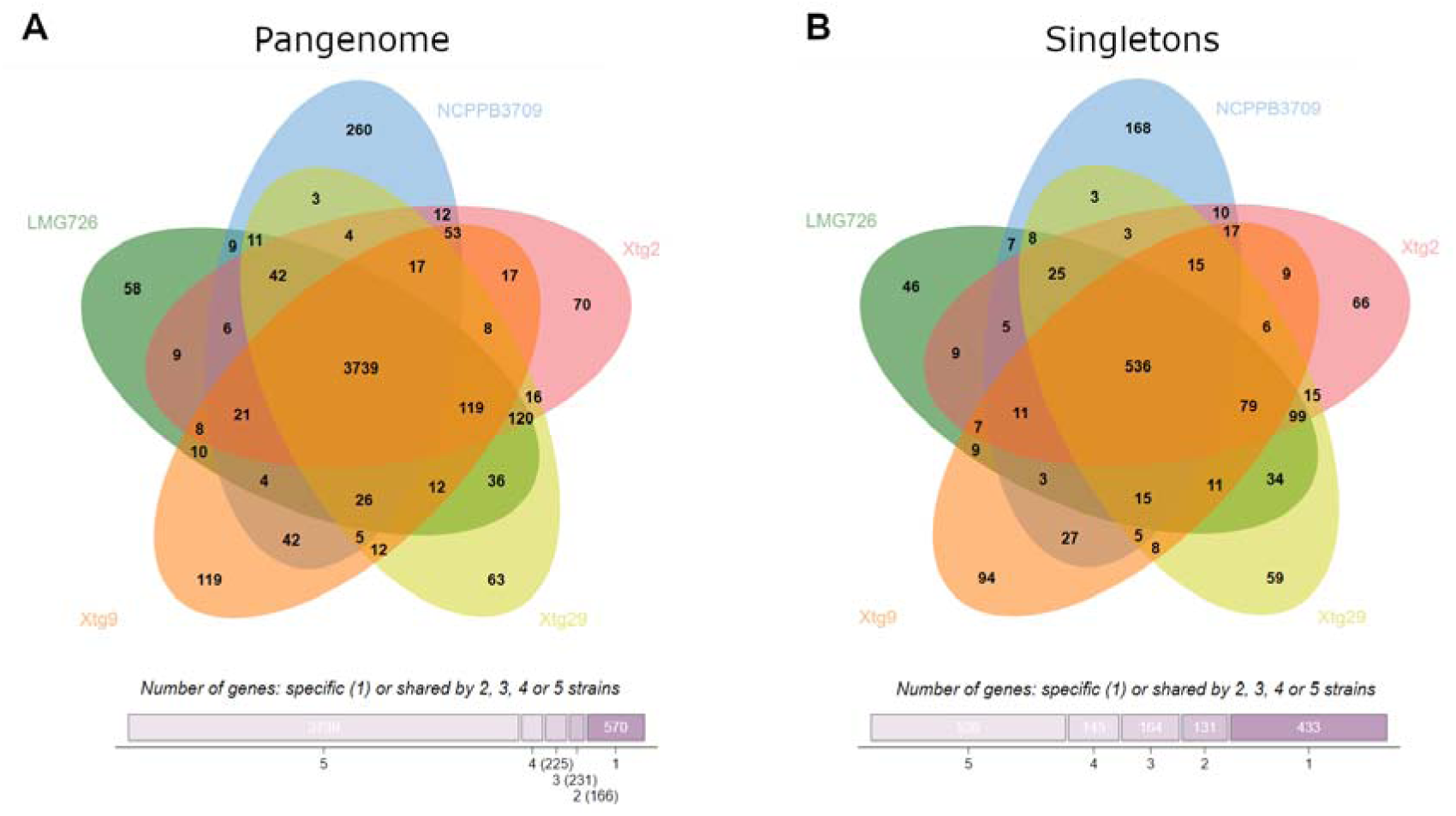
A) Pangenome (i.e., total gene content) of the five *Xanthomonas translucens* pv. *graminis* strains investigated, and B) content in singletons (i.e., genes that were exclusive to *X.t.* pv. *graminis* strains). Numbers indicate the number of genes shared by specific sets of strains. Bottom bars indicate the number of genes that shared by five, four, three or two strains or that are specific to one strain.

**Table 2.**
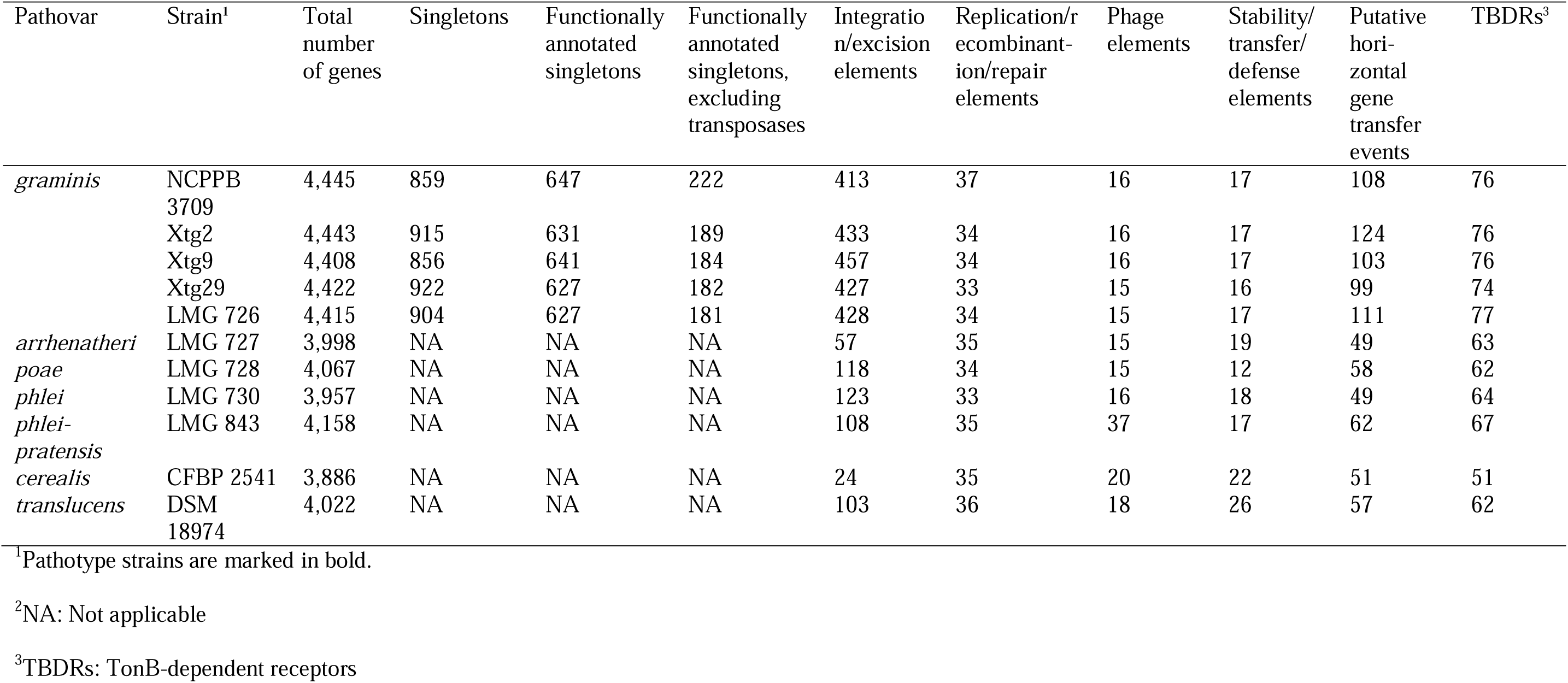
Gene content of the *Xanthomonas translucens* strains compared in this study.

The functional characterization based on COG (Fig. 4) showed that many genes within *Xtg* could not be functionally annotated (13-15%) or were of unknown function (15-16%). The largest category with a known function was replication, recombination, and repair, with 733 to 781 genes of this category per strain (16-18%). Furthermore, this category was also largely represented in singletons (48-53%), ranging from 424 singleton genes of this category in NCPPB 3709 to 453 in Xtg9. Investigation of these genes revealed that most of them were annotated as transposases, suggesting a high transposase activity in *Xtg*. Further investigation of mobile genetic elements in the *Xtg* genome sequences showed an overrepresentation of integration/excision elements, corresponding with the previously identified transposases, with 413 to 457 of such elements found per strain, compared to 24 to 123 in other strains of the species (Table 2). The number of other types of genetic mobile elements was similar between *Xtg* and other pathovars. Additionally, *Xtg* strains displayed a higher number of putative horizontal gene transfer events, with 99 to 124 predicted putative events, compared to 49 to 62 in the rest of the species (Table 2).

**Figure 4.**
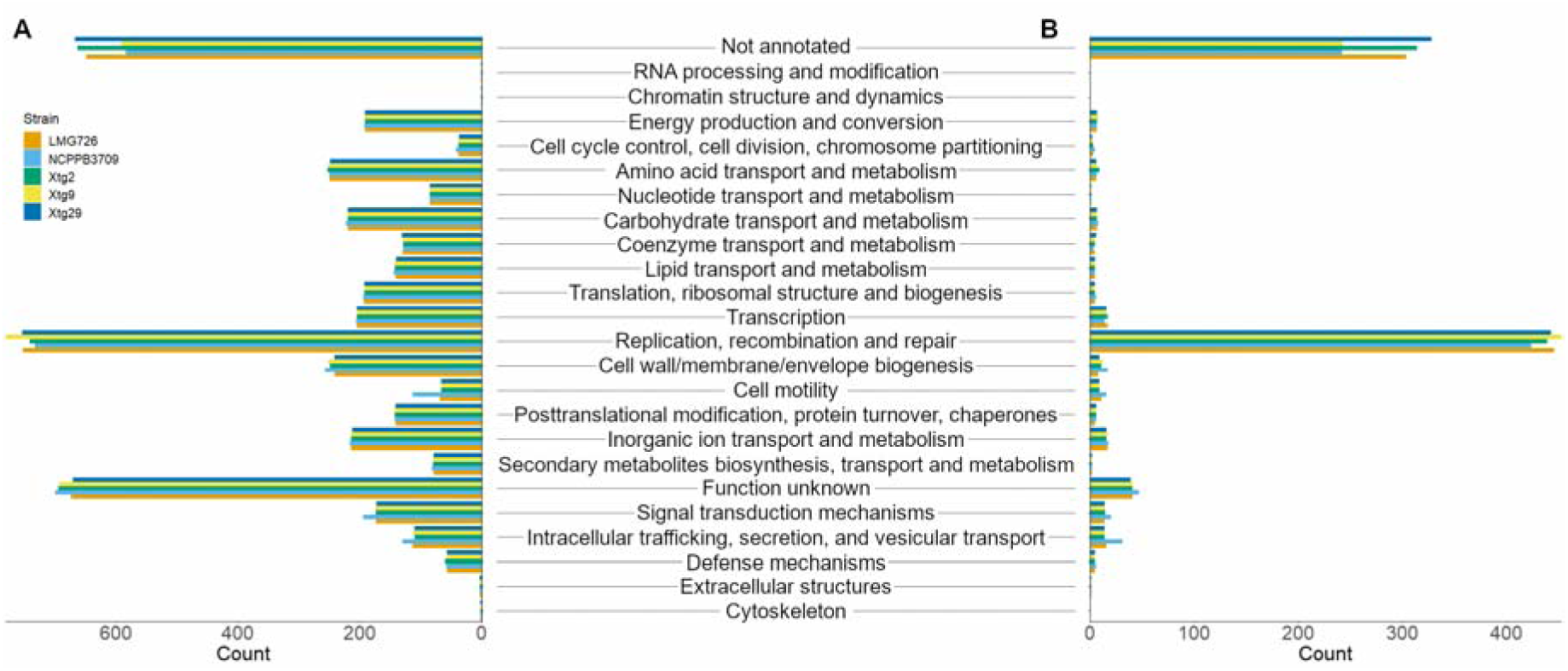
Clusters of orthologous groups (COG)-based functional characterization of A) the total gene content of the five *X. translucens* pv. *graminis* strains investigated, and B) singletons, genes that were exclusive to *X. t.* pv. *graminis* strains.

Investigation of secretion systems showed that all *Xtg* strains have the *xps* T2SS and the T3SS but not the *xcs* T2SS and T4SS (Fig. 5). Furthermore, a T6SS-i4 was found in NCPPB 3709, Xtg2 and Xtg9, but no T6SS could be found in LMG 726 and Xtg29. The type III effector (T3E) repertoire of *Xtg* was comparable among the strains of the pathovar, though only one effector of the XopP family was found in LMG 726 and Xtg29, compared to two in the other strains, and a XopL family effector was identified in LMG 726 only. When comparing to other pathovars, differences could be seen in the number of effectors of some families, such as AvrBs2 and XopF effectors, with only one found in *Xtg*, compared to two in most other pathovars, though only one XopF effector was found in LMG 730 and LMG 843 as well (Fig. 5). No XopG effector could be identified in *Xtg*, while all other pathovars possessed one. Moreover, aside from one gene found in LMG 726, no XopL effector was found in the other four *Xtg* strains, when other pathovars possessed one to three effectors of this family. While all other pathovars of clade Xt-III possessed exactly two XopX effectors, three could be found in *Xtg*, as was the case for pv. *translucens* (Xt-I) and pv. *cerealis* (Xt-II). Finally, three genes of the XopE family were identified in all *Xtg* strains, while only two were found in CFBP 2541, one in LMG 730, LMG 843 as well as DSM 18974, and none were found in LMG 728 and LMG 727. Overall, the *Xtg* strains displayed the smallest T3E set, ranging from 20 to 21 per strain, compared to 24 to 26 in other pathovars of clade Xt-III and 31 in pv. *translucens* and pv. *cerealis*. No TAL effector could be identified in any *Xtg* strain.

**Figure 5.**
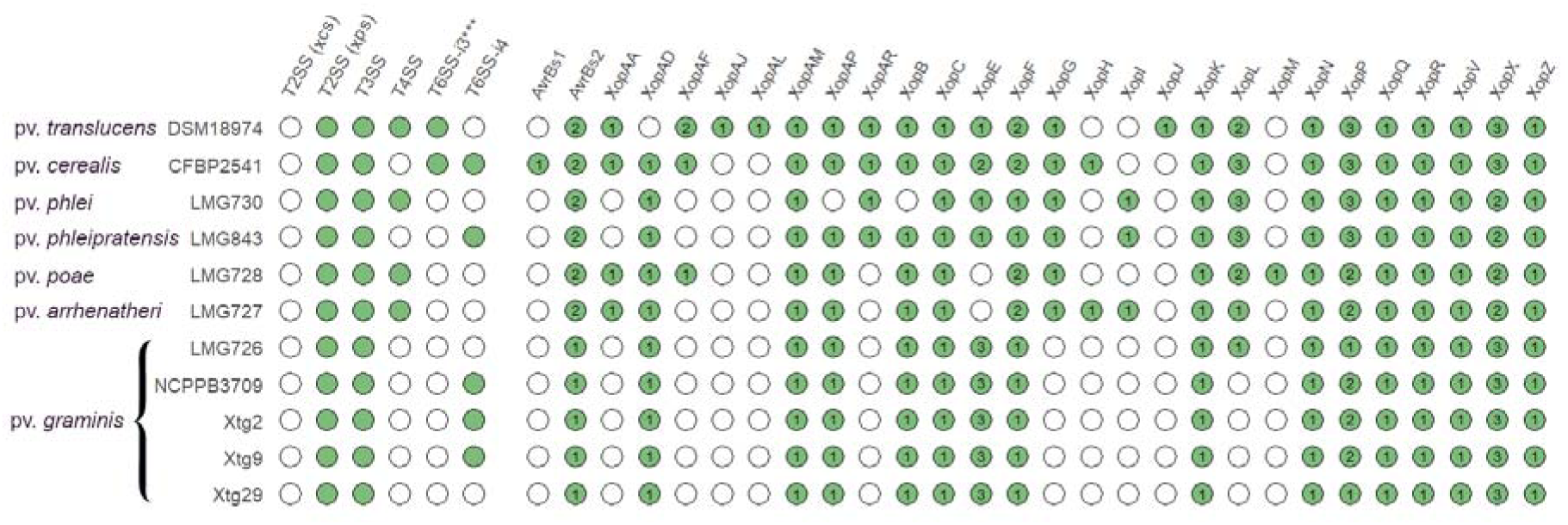
Presence of the *xps* and *xcs* type II secretion systems (T2SS), type III secretion system (T3SS), type IV secretion system (T4SS), subtype 3*** type VI secretion system (T6SS-i3***), and subtype 4 type VI secretion system (T6SS-i4) gene clusters, as well as putative type III effectors in the genome sequences of *Xanthomonas translucens* strains. Green circles indicate presence and white circles indicate absence. Numbers denote the number of putative effectors of each family.

Among the 856 to 922 singletons observed in *Xtg* strains, 627 to 647 were functionally annotated (Table 2). After excluding transposases, 181 to 222 annotated singletons could be identified, of which 148 were shared by all strains (Table 2). Among these singletons, genes that could have a role in the pathogenicity of *Xtg* were further investigated. As secreted or cell surface proteins are often involved in interaction with other organisms, we focused on genes that harbored a predicted secretion signal or a putative non-classical secretion. Among these, genes encoding for a filamentous hemagglutinin family outer membrane protein, a YadA family autotransporter adhesin, ten TonB-dependent receptors (TBDRs), and 11 degradative enzymes, including two subtilases, two esterases, a serine hydrolase, a glycerophosphodiester phosphodiesterase, a pectate lyase, a phospholipase, an alpha-galactosidase, a beta-galactosidase, and an aminopeptidase, were found. Seven additional TBDRs that were not shared by all strains were also found. Of the 17 identified TBDRs, five were annotated as diverse iron-siderophore receptors, five as oar-like proteins, three as vitamin B12 transporters, and one as a colicin receptor. However, while TBDRs were overrepresented in *Xtg*, ranging from 74 to 77 per strain, compared to 51 to 67 in other pathotype strains of the species (Table 2), many genes with similar annotations were also found among the TBDR genes that were conserved between pathovars, suggesting a functional redundancy of these TBDRs.

Additionally, a second gene predicted as encoding a filamentous hemagglutinin gene was found neighbouring the first one, being shared by all strains except NCPPB 3709. Further investigation of the locus containing the two genes revealed the presence of transposases in between them, suggesting it may be a single gene that was truncated by the insertion of these transposable elements. A BLASTn comparison with similar partial filamentous hemagglutinin genes that were previously identified in a draft genome sequence of Xtg29 revealed they corresponded to the same locus. One of the previously identified genes corresponded to the 5’-most gene that was identified in the singleton analysis, while the second gene corresponded to an additional smaller gene downstream of the two other genes we identified, which was present in all strains except NCPPB 3709 (Figure 6). A BLASTx comparison of the hypothetical full-length gene, consisting of the combined partial sequences, revealed a high identity to other filamentous hemagglutinin genes from *Xanthomonas* spp., further confirming this locus corresponds to a single gene that was truncated.

**Figure 6.**
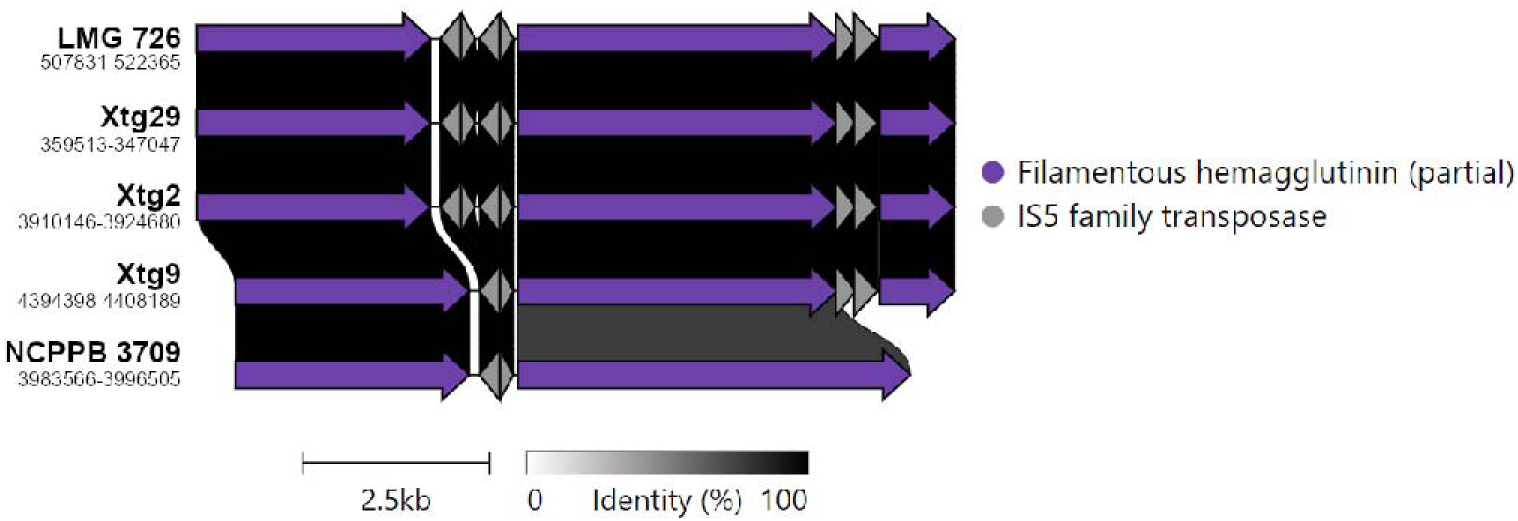
Pairwise comparisons of the filamentous hemagglutinin locus within *Xanthomonas translucens* pv. *graminis*. Colors denote similar genes. Numbers below strain names refer to the position of the gene cluster in this genome sequence.

Furthermore, a cluster containing five minor pilins of type IV pili (T4P) *fimT*, *pilV*, *pilW*, *pilX*, and *pilE*, as well as the anti-retraction factor *pilY1* was also found. This cluster is also present in other *X. translucens* pathovars, but the cluster found in *Xtg* is highly divergent from the one found in other pathovars, and these genes were thus defined as singletons by the roary pipeline (Figure 7A). An in-depth analysis of the minor pilins revealed that the *pilX* gene lacked the characteristic N-terminal methylation domain and class III signal peptidase cleavage site in any *X. translucens* strains investigated, suggesting it may not be functional. In contrast, these domains were present in *pilE*, *pilV* and *fimT* for all strains. When investigating *pilW*, the gene was predicted as encoding a putative pilin only in *Xtg*, pv. *phleipratensis* strain LMG 843 and pv. *poae* strain LMG 728 (Figure 7B). However, the conserved N-terminal phenylalanine residue was present only in LMG 843 and was replaced by a leucine in LMG 728 and all *Xtg* strains.

**Figure 7.**
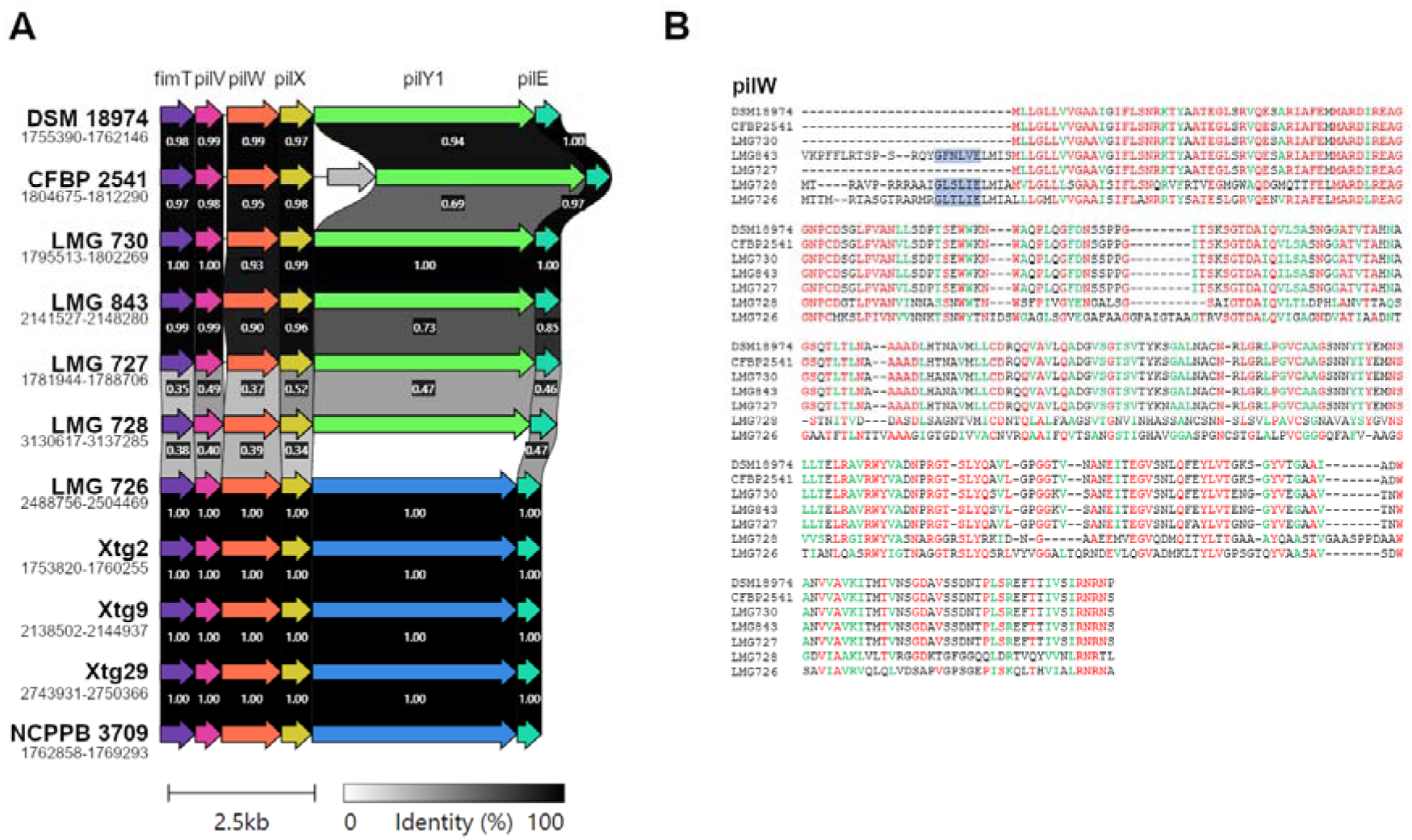
A) Pairwise comparisons of the minor pilins gene cluster between strains of *Xanthomonas translucens* spp. (i.e. *X.t.* pv. *translucens* DSM 18974, pv. *cerealis* CFBP 2541, pv. *phlei* LMG 730, pv. *phleipratensis* LMG 843, pv. *arrhenatheri* LMG 727, pv. *poae* LMG 728, and pv. *graminis* LMG 726, Xtg2, Xtg9, Xtg29 and NCPPB 3709). Colors denote similar genes. The *pilY1* gene is represented in different colours in pv. *graminis* strains compared to other strains as its sequence was too divergent and they were considered as different genes by the software. Numbers below strain names refer to the position of the gene cluster in this genome sequence. B) Alignment of the amino acid sequences of pilW. Only the pv. *graminis* strain LMG 726 is represented as the sequences were identical in all five pv. *graminis* strains. Colours represent amino acids that were conserved among at least six out of seven strains (red) or amino acids with similar properties that were found in at least six out of seven strains (green). Highlighted in blue is the class III signal peptidase cleavage site that is characteristic to pilins.

## 4 Discussion

Comparative analysis of five *Xtg* strains revealed a very high genome plasticity within the pathovar compared to all other pathovars of *X. translucens*. However, while the strains showed 99.9 to 100% nucleotide identity, the analysis of conserved genomic regions showed a high level of rearrangement between the five strains with 65 to 125 LCBs. This is the first report of such a high level of rearrangements within *Xtg*, which could be observed thanks to the generation of complete genome sequences that allowed a clear observation of the genome organization in each strain. In comparison, a similar analysis with data published by Shah et al. (44) and using complete genome sequences of four pv. *translucens* and eight pv. *undulosa* strains with ANI values as low as 98.9 and 99.5%, respectively, showed a considerably lower amount of rearrangements, with 53 to 67 and 39 to 67 LCB (38, data not shown). This is in line with a previous comparative analysis within the *X. translucens* species that showed that clade Xt-III, which contains *Xtg*, was the clade showing the most rearrangements (3). Additional genome sequences for other pathovars of clade Xt-III would help to determine whether this high level of plasticity is exclusive to *Xtg* or whether it is a characteristic of the entire clade.

The singleton analysis revealed 1,409 genes that were exclusive to the *Xtg* strains, with 536 of them being found in all five strains. Functional characterization revealed that most of these singletons were annotated as transposases, and further analysis of mobile genetic elements also revealed a high amount of insertion/excision elements in the genomes of *Xtg* strains, corresponding to the identified transposases. Such a high number of transposable elements in *Xtg* could explain the high genomic plasticity that was observed, as they are genetic elements that are mobile within the genome and can facilitate gene duplications, chromosomal recombinations and horizontal transfer (48,49). A high genome plasticity has previously been found in many *Xanthomonas* spp., and was suggested to play an important role in host adaptation (50). A comparably large number of insertion sequences was reported in *X. oryzae* pv. *oryzae*, where these numerous mobile elements were suggested to play a role in race differentiation (51). Insertion elements can also cause the loss of gene function, and virulence can emerge from the inactivation of avirulence genes (52,53). Furthermore, transposable elements have been suggested to play a role in the movement of “passenger genes” in *Xanthomonas* spp., including virulence genes, contributing to their diversification and spread (54,55). As such, mobile genetic elements can play a crucial role in pathogen adaptation, and the large amount of such elements in *Xtg* compared to other pathovars of the species could, at least partially, explain its high virulence and broad host range.

The total number of genes found in each strain of the pathovar was comparable, confirming the high number of genes already identified in the *Xtg* pathotype strain LMG 726. When compared with other pathotype strains, LMG 726 exhibited the highest number of genes of the species, as well as the highest number of genes that were exclusive to the strain (3). While in this previous comparative analysis, as much as 950 genes were found to be exclusive to LMG 726, this number was influenced by the strains included in the comparison. The finding of 1,409 genes exclusive to *Xtg*, with 536 shared by all strains, using all available complete genome sequences of *X. translucens* strains is most likely a better estimation of the number of genes specific to *Xtg*. However, this unusually high amount seems to mainly be explained by the abundance of transposases in the pathovar. Nonetheless, many singletons that could play a role in the pathogenicity and host range of *Xtg* have been uncovered.

First, a cluster containing five minor pilins and the anti-retraction factor *pilY1* was found to greatly differ between *Xtg* and other pathovars of the species. Thesee pilins are components of T4P, which mediate surface adhesion, biofilm formation and twitching motility (56,57). T4P consist of a filamentous polymer of the major pilin PilA, and minor pilins may be incorporated into the pilus. Further investigation of minor pilins of this cluster revealed that only *pilE*, *pilV* and *fimT* may be functional in all pathovars of the species. For *pilW*, the N-terminal cleavage and methylation domain that is characteristic to pilins and required for their processing was present only in pvs. *graminis*, *poae*, and *phleipratensis*, though the conserved phenylalanine that undergoes methylation after cleavage was only present in pv. *phleipratensis* and was substituted by a leucine in the other two pathovars. However, it was previously shown that although the phenylalanine residue is highly conserved in pilins, its substitution by another amino acid did not affect secretion, cleavage and methylation of the pilin and subsequent T4P assembly (58). It is thus plausible that *pilW* is still functional in pvs. *graminis* and *poae*. The six identified genes, as well as the major pilin PilA were previously shown to differ significantly in Xtg29 compared to strains of other *X. translucens* pathovars, while the genes encoding the main components of the T4P machinery were highly conserved (17). That this divergent cluster is conserved within *Xtg* could indicate a role in the virulence of the pathovar. Indeed, the T4P is an extracellular feature of the bacteria that may be detected by plant defense mechanisms. The high sequence variation in the minor pilins, as well as the major pilin PilA may allow *Xtg* to avoid plant defenses while mediating surface adhesion and biofilm formation.

Among other singleton genes, two filamentous hemagglutinin genes were identified. However, further investigation revealed that they likely correspond to a single gene in which multiple transposases were inserted. Moreover, these genes corresponded to the two partial filamentous hemagglutinin genes that were previously reported as singletons of *Xtg* (17). The N-terminus of filamentous hemagglutinins possesses a signal peptide, as well as a two partner system domain that are involved in their secretion to the periplasm and the apoplasm, respectively (59,60). Whether the gene encoding for the N-terminal part of the protein is still functional remains to be determined, though it is unlikely, as essential domains may be found in the C-terminal part of the gene. In *X. axonopodis* pv. *citri*, filamentous hemagglutinins were shown to be crucial to virulence by promoting plant surface adhesion and biofilm formation but acted as elicitors of plant defense in *X. campestris* pv. *vesicatoria* (61,62). Thus, the loss or truncation of this gene in *Xtg* may help it evade recognition by the plant host.

Additionally, 17 singleton genes coding for TBDRs were found, with ten being shared by all *Xtg* strains. TBDRs are known for their role in iron and vitamin B12 uptake and are mostly regulated by the ferric uptake regulator gene (63). Mutants of this regulator were unable to induce disease symptoms in *X. o.* pv. *oryzae* which was hypothesized to be due to an inability to cope with the oxidative stress conditions encountered during the infection, emphasizing the role of TBDRs in virulence (64). Moreover, in *X campestris* pv. *campestris*, some TBDRs were found to be activated by HrpG and HrpX, regulatory proteins of the type III secretion system, and the SuxA TBDR was found to play a major role in pathogenicity by acting on sucrose import (65). A comparative analysis across 226 eubacterial genome sequences showed that TBDRs were overrepresented in *Xanthomonas* spp. compared to other bacteria, ranging from 36 TBDRs in *X. oryzae* pv. *oryzae* to 68 in *X. axonopodis* pv. *citri* (65). In this study, similar numbers of TBDRs were found in all strains of *X. translucens*, with an overrepresentation in *Xtg*. The identified singletons were mainly annotated with functions related to iron and vitamin B12 uptake, and similar genes were found to be shared across the species. It is thus unclear if these 17 singletons serve a specific function in the pathogenicity of *Xtg*, or if they are functionally redundant with other TBDRs.

A total of 11 degradative enzymes were also found in singletons that were shared by all *Xtg* strains and were predicted to be secreted. Such enzymes are often secreted by the T2SS and involved in the degradation of the plant cell wall (13,15). These include cellulases, xylanases, pectate lyases, and polygalacturonases, which degrade the main polysaccharide cell wall constituents, as well as various proteases and lipases. *X. oryzae* pv. *oryzae* mutants lacking a T2SS were shown to be virulence deficient and multiple enzymes secreted by the T2SS were shown to be necessary for virulence (66–68). As *Xtg* mutants lacking a T3SS showed a drastic reduction in symptoms, but still survived in the plant, it was hypothesized that other virulence factors may be important in *Xtg* (12). Therefore, the T2SS and the enzymes it secretes may play a crucial role in the *in planta* growth of *Xtg*.

Finally, a gene encoding for a YadA family autotransporter adhesin was identified as a singleton shared by all *Xtg* strains. Such adhesins showing a high similarity to YadA from *Yersinia* spp. have been shown to mediate cell to cell aggregation, biofilm formation, and adhesion to the host cell surface (69,70). In *Xanthomonas oryzae* pv. *oryzae*, mutations in the genes encoding for the YadA-like adhesins XadA and XadB were shown to result in reduced surface attachment and virulence after surface inoculation (71,72). In *Xtg*, this adhesin may therefore play a similar role in attachment to the host.

An in-depth analysis of the secretion systems within *Xtg* showed that a T6SS-i4 was present in strains Xtg2, Xtg9 and NCPPB 3709 and not in LMG 726 and Xtg29, confirming previous reports investigating these strains (17). It was previously hypothesized that the T4SS and T6SS could act as elicitors of plant defense, and that their absence in LMG 726 could help it evade plant defenses (3). However, the presence of a T6SS in Xtg2, Xtg9 and NCPPB 3709 may disprove this hypothesis. Nonetheless, its absence in Xtg29 and LMG 726 indicates that it is not crucial to the pathogen’s survival and virulence. Conversely, the *xps* T2SS and the T3SS were found to be conserved, suggesting they may be crucial to the pathovar.

The T3E repertoire within the pathovar was largely conserved, except for only one XopP family effector being present in strains Xtg2, Xtg9 and NCPPB 3709, compared to two in LMG 726 and Xtg29, as well as one XopL family effector being present only in the LMG 726 strain. This XopL effector could have been conserved in LMG 726, or it might have been recently acquired through horizontal transfer. Nonetheless, as it is unique to LMG 726, it is likely not crucial to the lifecycle of *Xtg*. Rather, the absence of XopL effectors may have been beneficial to other strains of *Xtg*, as they all lacked such effectors, while all other pathovars of the species possessed at least one and up to three. Similarly, several effectors appear to have been lost in *Xtg*, such as AvrBs2, XopF, or XopG, as the pathovar possesses fewer of these effectors than other pathovars. Furthermore, a comparable loss of a XopP effector may have occurred in strains LMG 726 and Xtg29. Interestingly, *Xtg* possesses the smallest set of T3E in the species, which may be explained by such a loss of effectors during evolution. Finally, no TALE effectors were identified, further confirming the absence of such effectors in the pathovar (3,12). This suggests that these T3E were either unnecessary or even deleterious to the pathovar. As these effectors could trigger plant defence, their loss may have contributed to the success of *Xtg* as a pathogen of many grass species. On the other hand, three effectors of the XopE family were present in *Xtg*, while only two, one, or none were found in other pathovars. Additionally, three XopX effectors were found in *Xtg*, while other pathovars of clade Xt-III possessed only two. XopE effectors belong to the HopX family, which contains transglutaminases and possess a catalytic site that may be involved in proteolysis (73). The XopE2 effector was shown to suppress hypersensitive response in *Nicotiana* spp. and to inhibit pattern-triggered immunity (74,75). XopX effectors were found to be required for virulence in *X. euvesicatoria* on tomato and pepper by interfering with plant immune responses (76,77). This suggests that XopE and XopX effectors may play a role in the pathogenicity of *Xtg* by suppressing plant defences.

Phylogeny based on ANI showed that all five available *Xtg* strains were highly identical, with only the Norwegian NCPPB 3709 strain being showing some differences to the other *Xtg* (99.9% ANI). A previous analysis of the genetic diversity of *Xtg* using AFLP markers also showed that multiple strains from Switzerland, France and Belgium displayed a very low diversity (40). In other pathovars of the species, a higher diversity was observed, with only 99% ANI within pv *undulosa* and pv. *Translucens* (44). Although these strains were from a more diverse origin than in *Xtg*, with strains from North and South America as well as Iran, there was still a high diversity when comparing strains of a similar origin. This further highlights the particularity of the low diversity observed within *Xtg*. Still, as aside from one Norwegian strain, all *Xtg* strains compared in this study originate from Switzerland, obtaining additional complete genome sequences from strains of more diverse origins would help to better determine the genomic diversity existing within the pathovar.

Overall, many common features of *X. translucens* were lacking in *Xtg* and seem to have been lost over the course of evolution, as they were present in the rest of the species. No T4SS was found in the pathovar and a T6SS was found only in strains NCPPB 3709, Xtg2 and Xtg9. It had the smallest set of T3E and no TALE were identified. Previous research also showed that although a nearly complete flagellar gene cluster is found in strain NCPPB 3709, such a cluster was absent in all other strains investigated (17). In this regard, the high mobile genetic element activity observed in *Xtg* could have played an important role in this loss of virulence features through transposable element insertion within a gene, or by facilitating chromosomal region deletions (52,53,78). Furthermore, a highly divergent T4P was found, both in terms of the amino acid sequence of the pilins, as well as in the presence or absence of functional minor pilins, compared to the rest of the species. This loss or divergence of main virulence features in *Xtg* may be responsible for its ability to infect a large range of plant hosts, by allowing the bacteria to evade plant defense mechanisms.

## 5 Conclusion

Our study provided new high-quality complete genome sequences for *X. translucens Xtg*, allowing for an in-depth comparative genome analysis within the pathovar. This revealed that *Xtg* exhibits a remarkable genome plasticity, likely related to an unusually high amount of transposases, which could explain its success as a critical pathogen of a large range of forage grasses. A set of potential virulence factors of the pathovar were identified, and a comprehensive analysis of these genes will contribute to better understand its virulence and host range, providing a basis for the development of new cultivars of forage grasses with increased resistance to the disease.

## 6 Declarations

### 6.1 Competing interests

The authors declare that the research was conducted in the absence of any commercial or financial relationships that could be construed as a potential conflict of interest.

### 6.2 Funding

Funding was provided by the Swiss National Science Foundation (Grant No: IZCOZO_177062).

### 6.3 Authors’ Contributions

Design of the study: FG and RoK, with the help of BS. Genome sequencing, assembly and data analysis: FG with the help of VR and RaK. Data interpretation, writing and reviewing: FG, RoK, VR, RaK and BS. All authors contributed to the article and approved the submitted version.

## Acknowledgments

This article is based upon work from COST Action CA16107 EuroXanth, supported by COST (European Cooperation in Science and Technology). Library preparation and sequencing were done by the Functional Genomics Center Zurich (Zurich, Switzerland). RaK and RoK are grateful for an international mobility EXPLORE grant from the Montpellier University of Excellence (MUSE).

## 6.4.1 Availability of data and materials

The genome sequences generated for this study can be found in the NCBI GenBank repository under the accession numbers specified in Table 1.

## 6.5 Ethics approval and consent to participate

Not applicable

## 6.6 Consent for publication

Not applicable

## Notes

### Competing Interest Statement

The authors have declared no competing interest.

